# Phase transition of human gastrointestinal mucin solutions

**DOI:** 10.1101/2025.10.21.683622

**Authors:** Yi Hui Zhao, Li Ming Zhang, Di Jia

## Abstract

In this study, we investigate the phase behavior of mucin aggregates from gastrointestinal mucus in the addition of multivalent salts. A reentrant phase transition is observed with trivalent and tetravalent salts. Mucin aggregates exhibit liquid-solid phase separation in divalent salts such as Ca^2+^, Cu^2+^ and Ba^2+^, whereas the solution remains homogeneous across all tested concentrations of MgCl_2_. The ability of multivalent salts in promoting protein participates follows the order: spermine>Fe^3+^>Al^3+^>Ba^2+^>Cu^2+^>Ca^2+^>Mg^2+^, which is different from the Hofmeister series. What’s more, precipitates formed in CaCl_2_solution can be redissolved through ion exchange by adding NaCl or MgCl_2_.

## Introduction

The gastrointestinal mucus barrier is a crucial component of the human innate defense system, playing a vital role in regulating gastrointestinal microecology and maintaining gastrointestinal homeostasis. The major building blocks in mucus are mucins, which are large, highly glycosylated proteins and playing a pivotal role in ensuring the lubrication properties and barrier function of mucus (1). The mucins are >80% carbohydrate, and are concentrated into mucin domains (2). These domains are built on a protein core that is rich in the amino acids proline, serine and threonine (called PTS sequences) (3). Mucins feature cysteine-rich regions at both the amino and carboxy termini, Mucin-2 can form an extensive net-like polymer structure, where the C-termini of individual Mucin-2 molecules create dimeric complexes through disulfide bonds, while the N-termini form trimeric complexes (1).

Ions play a key role for many biological processes on the molecular scale, such as molecular interactions, self-organization and assembly, reaction equilibria and recognition *etc*.(4). For example, the Mg^2+^ are required to screen the highly negative charge of nucleic acids to enable the nucleic acid-protein interactions (5). High Ca^2+^ ion concentration alongside low pH enables mucus packing by masking negatively charged glycans on the Mucin-2 protein, which is the component of mucus in intestine (6, 7). For gastrointestinal mucin proteins with charged amino acid, the effect and phenomena induced by multivalent ions are complex and it is important to understand the interaction between mucin and multivalent ions and the consequences of this ion binding such as collapse, charge reversal, precipitation and reentrant of the mucin.

In this study, we flashed the gastrointestinal mucus by a large amount of normal saline, after centrifugation and filtration with a filter membrane, the mucin aggregates can be obtained. The phase behavior of mucin aggregates from gastrointestinal mucus in the addition of multivalent salts was investigated. A reentrant phase behavior of mucin aggregates can be observed in trivalent salts. The mucin aggregates showed liquid-solid phase separation behavior in divalent salts such as Ca^2+^, Cu^2+^ and Ba^2+^, while the mucin aggregates solution remains in one-phase for MgCl_2_ solution in the entire range of measurement. What’s more, by adding MgCl_2_ solution in the precipitate of CaCl_2_, the precipitate redissolved in MgCl_2_ solution through ion-exchange. We found that the ability of multivalent salts is Fe^3+^>Al^3+^>Ba^2+^>Cu^2+^>Ca^2+^>Mg^2+^, which is different from the Hofmeister series (8). The complete phase behavior with multivalent ions has been explained in terms of the interplay of electrostatic repulsion and ion induced short-range attraction between the mucin aggregate.

## Results and discussion

The gastrointestinal mucus was flushed with a large volume of normal saline. Following centrifugation and filtration through a filter membrane, mucin aggregates were obtained. The hydrodynamic radius (R_h_) of the mucin aggregates is precisely quantified by using dynamic light scattering (DLS). The DLS results indicates a monodisperse size distribution with an average R_h_=116nm (Fig. 1a, b) (9), suggesting a homogeneous self-assembled structure. Various types and concentrations of pre-filtered salt solutions were then added to the mucin aggregate solutions.

**Fig. 1.**
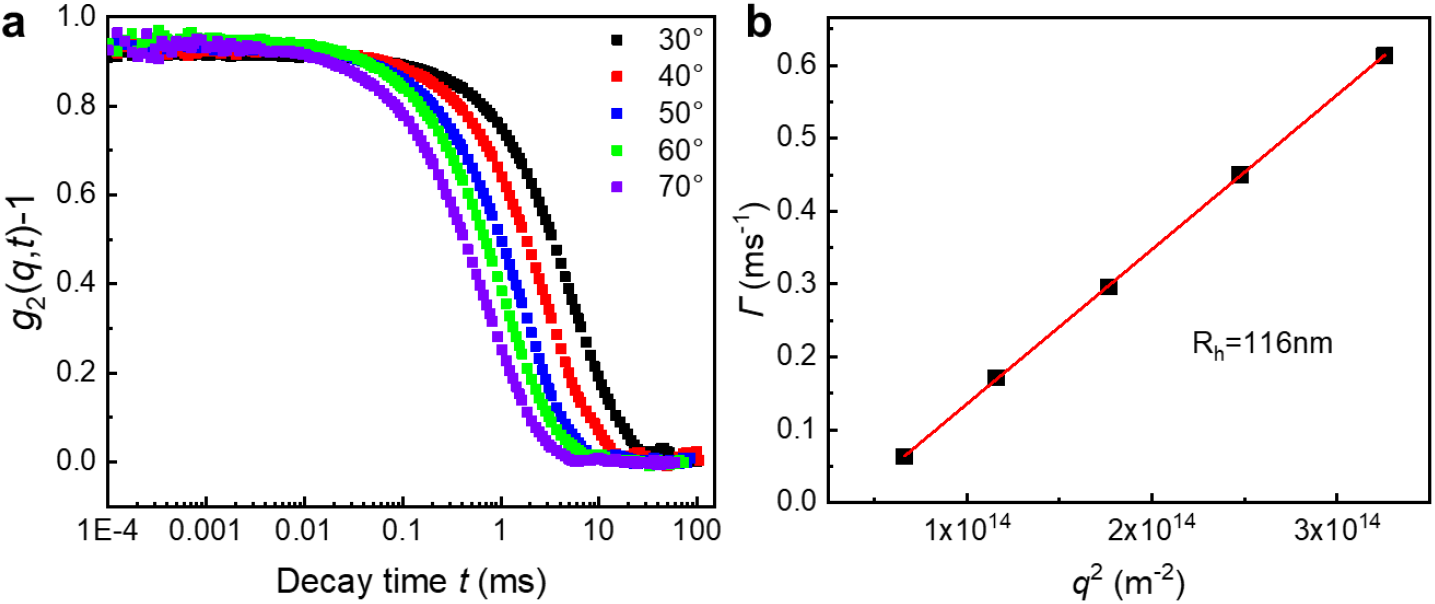
(a) DLS data of the native mucin aggregate solution. (b) *q*^2^ dependence of the relaxation rate *Γ* for the native mucin aggregate solution.

Figure 2 illustrates the phase behavior of mucin aggregate solutions in the presence of trivalent salts. As shown in Fig. 2a, at low concentration of AlCl_3_, the system remains stable as a one-phase mucin solution. With increasing AlCl_3_ concentration, liquid-solid phase separation occurs, resulting in the formation of a highly concentrated precipitate and a very dilute phase. The phase separation is driven by the electrostatic condensation of counterions along the highly charged mucin chains, the long-range Coulombic repulsion as well as the short-range bridging attraction (10). At even higher AlCl_3_ concentrations, a reentrant phase behavior of mucin aggregates is observed, wherein the precipitated mucin aggregates redissolve (Fig. 2a). The reversion to one-phase is attributed to the screening of the short-range electrostatic attractions between the monomers induced by ion condensation at elevated salt levels (11).

**Fig. 2.**
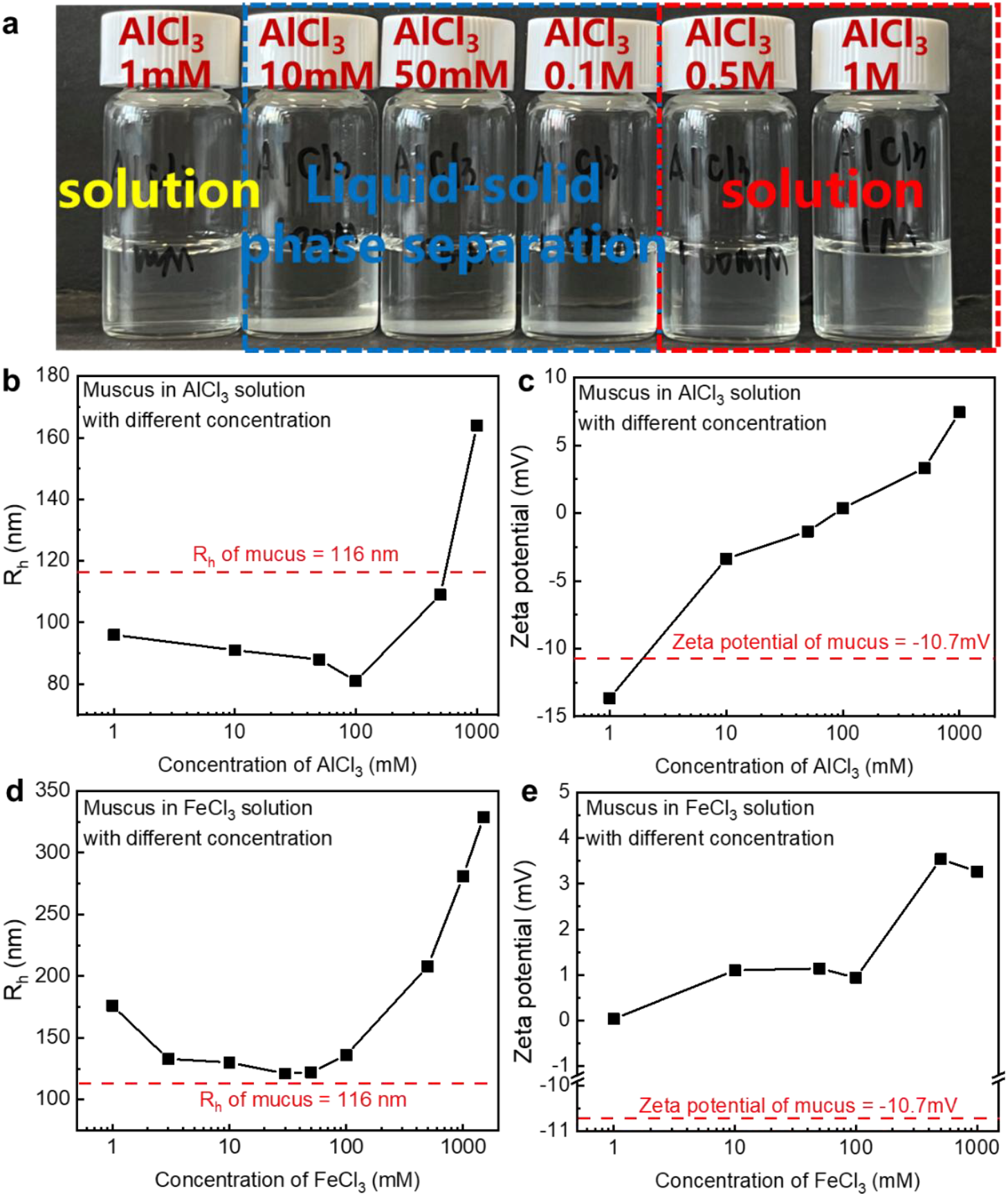
(a) Physical appearance of the mucin aggregates with varying concentration of AlCl_3_ salt. (b) Hydrodynamic radius (R_h_) of mucin aggregates as a function of concentration for AlCl_3_. (c) Zeta potential of mucin aggregate with varying concentration of AlCl_3_ salt. (d) Hydrodynamic radius (R_h_) of mucin aggregates as a function of concentration for FeCl_3_. (e) Zeta potential of mucin aggregate with varying concentration of FeCl_3_ salt.

The size of mucin aggregate in the supernatant was measured by DLS, as shown in Fig. 2b. Fig. 2c presents the corresponding zeta potential values of mucin aggregates across varying concentrations of AlCl_3_. With increasing concentration of AlCl_3_, the R_h_ of the mucin aggregates decreases within the liquid-phase separation region, while the zeta potential increases from approximately -11 mV to nearly 0 mV. This behavior can be attributed to the collapse of initially stretched mucin chains into compact, almost neutral structures. The collapse is driven by trivalent counterions that adsorb to the mucin backbone and act as bridges, enhancing intrachain attraction (12). When the AlCl_3_ concentration exceeds the charge reversal point (around 100mM), the system undergoes a phase transition from liquid-solid phase separation to a clear solution, accompanied by an increase in mucin aggregate size. Beyond this point, further addition of AlCl_3_ leads to condensation of trivalent counterions on the mucin backbone, resulting in effective charge reversal. Above the isoelectric point, the mucin aggregates reswell due to the repulsive interactions among the overcharged condensed trivalent ions (13, 14).

For mucin aggregates in the presence of trivalent Fe^3+^, the phase behavior is consistent with that observed in AlCl_3_ solutions, exhibiting a phase transition from a one phase solution to liquid-solid phase separation (Fig. 2d). Followed by reentrant dissolution into a homogeneous solution. Within the phase-separated region, the size of the mucin aggregates in the supernatant decreases slightly. The charge reversal point of mucin aggregates occurs at 1mM FeCl_3_, indicating that the charges on mucin aggregates are completely screened at 1mM FeCl_3_ (Fig. 2e), so that the is always larger than that of native mucin aggregates. Beyond the reentrance point, the aggregate size increases with further increasing concentration of FeCl_3_. Reentrant dissolution is observed at 50mM FeCl_3_, indicating a stronger ability of Fe^3+^ to promote mucin aggregate precipitation compared to Al^3+^.

The phase behavior of mucin aggregates in the presence of divalent salts was investigated. Liquid-solid phase separation occurs in solutions of BaCl_2_, CuCl_2_ and CaCl_2_, with no reentrant behavior observed within the range of salt concentrations studied. The minimum salt concentrations required to induce mucin aggregate precipitation is 1mM for BaCl_2_, 1.5mM for CuCl_2_ and 2mM for CaCl_2_, indicating that the ability to promote mucin aggregate precipitation follows the order: Ba^2+^>Cu^2+^>Ca^2+^.

With increasing salt concentration, the size of mucin aggregates remains largely unchanged in BaCl_2_, but decreases in both CuCl_2_ and CaCl_2_ due to ion bridging effects (Fig. 3a, b). In contrast, mucin aggregates remain one phase in MgCl_2_ solutions, with aggregate sizes similar to those in native state (Fig. 3c), suggesting that Mg^2+^ has the lowest propensity to promote mucin aggregate precipitation. Thus, the overall trend observed Ba^2+^>Cu^2+^>Ca^2+^>Mg^2+^ deviates from the classical Hofmeister series (8). The behavior of mucin aggregates in different salt can be attributed to electrostatic interaction between divalent cations and specific binding sites along the mucin backbone. The observed differences in precipitation efficacy likely arise from variations in the microstructure of the mucin protein and the spatial distribution of charges along the aggregate surface (15).

**Fig. 3.**
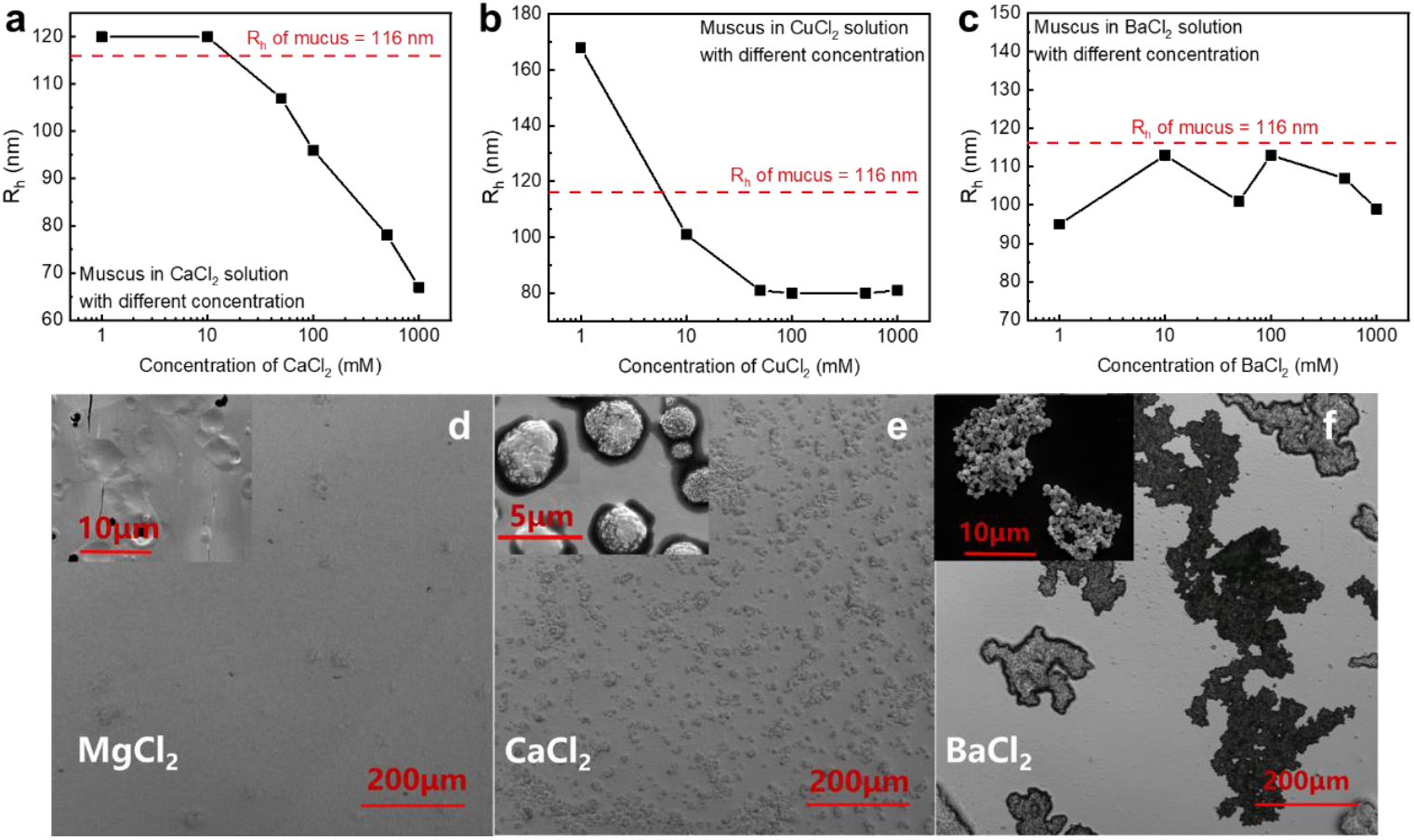
(a-c) Hydrodynamic radius (R_h_) of mucin aggregates as a function of concentration for CaCl_2_ (a), CuCl_2_ (b) and BaCl_2_ (c). (d-f) Optical microscope images of mucin aggregates in (d) 50mM MgCl_2_, (e) 50mM CaCl_2_ and (f) 50M MgCl_2_. Scale bars: 200μm (d-f). Inset show corresponding SEM images. Scale bars:10μm (d), 5μm (e) and 10μm (f).

Optical microscope images and SEM images show that the structural features and distribution of mucin aggregates precipitated in 50Mm MgCl_2_, CaCl_2_ and BaCl_2_. Almost no precipitate is observed in the presence of 50mM MgCl_2_, indicating that the mucin aggregate remains in one phase under this condition (Fig. 3d). In contrast, mucin aggregates form uniform homogeneously distributed spherical precipitates approximately 5μm in diameter in the presence of 50mM CaCl_2_, suggesting that Ca^2+^ promotes inter-aggregate linking into regular structures (Fig. 3e). While in 50mM BaCl_2_, mucin aggregates undergo large-scale precipitation (Fig. 3f). From a more microscopic perspective, individual aggregates appear collapsed into spherical units that are bridged by Ba^2+^, resulting in precipitates with irregular shapes and heterogeneous sizes. The imaging results are consistent with the trend in precipitation ability: no precipitate forms in MgCl_2_, well-distributed spherical precipitates form in CaCl_2_, and large, irregular patches form in BaCl_2_. The size and extent of the precipitation increase with the enhancing ability of the divalent salt to promote aggregation.

The effect of NaCl addition on mucin precipitates formed in 100mM CaCl_2_ is shown in Fig. 4a, c. After the precipitate forms, the supernatant is removed and replaces with an equal volume of NaCl solution at varying concentrations. As the concentration of NaCl increases, the precipitates initially formed in CaCl_2_ redissolve, which can be attributed to competitive binding between Ca^2+^ and Na^+^ counterions. Although Na^+^ has a lower binding constant than Ca^2+^, a large excess of Na^+^ can displace Ca^2+^ from the mucin aggregate precipitates, thereby reducing the number of condensed Ca^2+^. Consequently, when the concentration of Na^+^ exceeds that of Ca^2+^, the added NaCl screens the bridging attractions, leading to the redissolution of the precipitate and a transition from two phases to a single phase (11, 15). Furthermore, increasing the NaCl concentration leads to larger mucin aggregates, indicating that electrostatic screening promotes the redissolution of mucin aggregates.

**Fig. 4.**
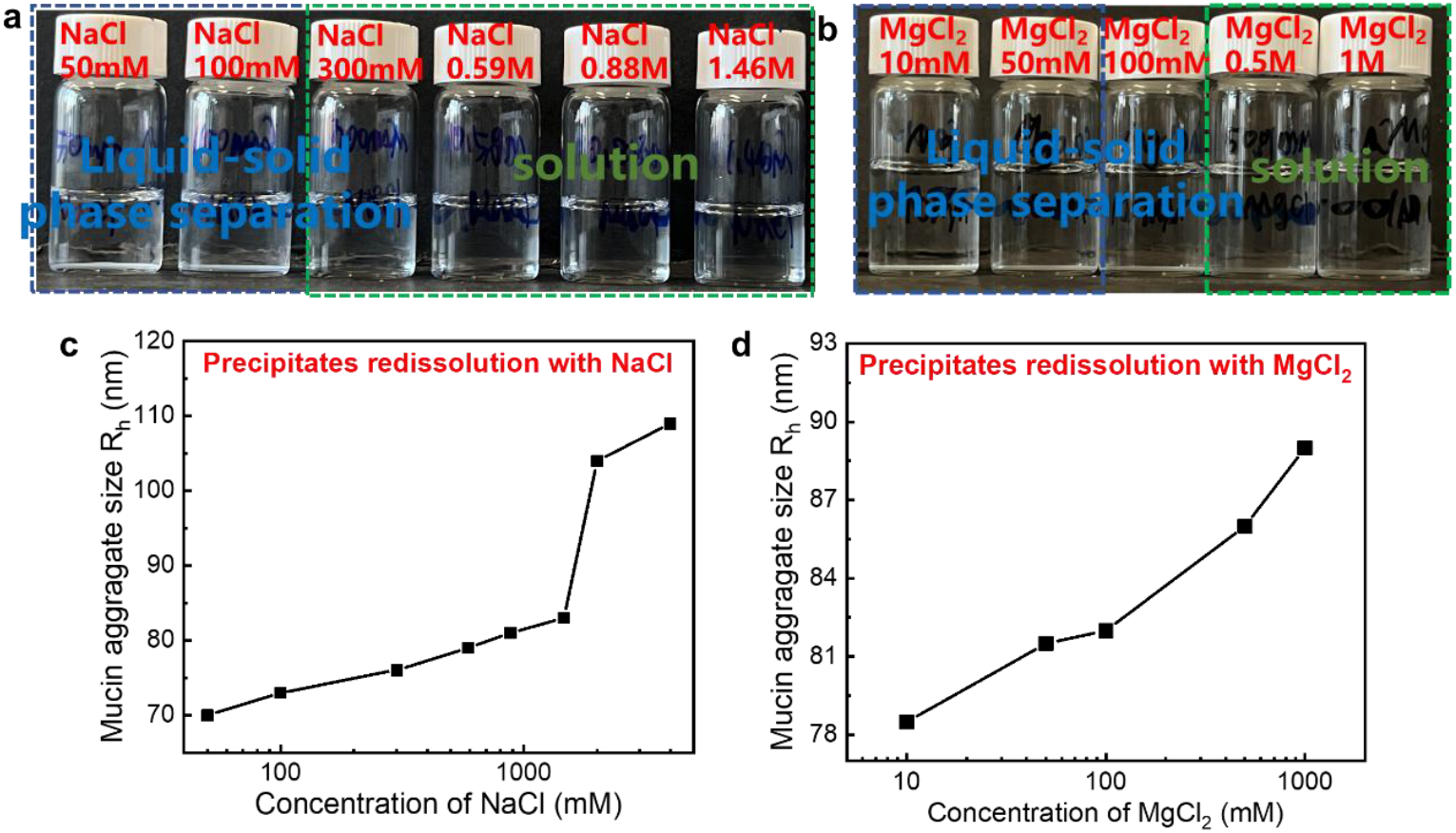
(a) Redissolution behavior of precipitates formed in 100mM CaCl_2_ upon addition of NaCl. (b) Redissolution behavior of precipitates formed in 100mM CaCl_2_ upon addition of MgCl_2_. (c) Hydrodynamic radius (R_h_) of mucin aggregates as a function of concentration for NaCl. (d) Hydrodynamic radius (R_h_) of mucin aggregates as a function of concentration for MgCl_2_.

A similar trend is observed when MgCl_2_ is added to mucin precipitates formed in 100mM CaCl_2_ (Fig. 4b, d). To compare the screening effects of monovalent and divalent salts on Ca^2+^, we calculated the Debye length (*ξ*) of the salt solution. For monovalent salts, Debye length is given by (16)

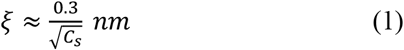

where C_s_ is the concentration in units of moles per liter. For divalent salts, Debye length is given by

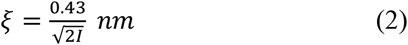

where *I* is the ionic strength, which is given by

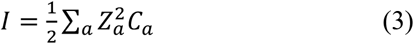

Z_a_ is the charges of ions and C_a_ is the concentration of ions of a-type in molarity. At comparable Debye lengths (*ξ*) – achieved with 200mM MgCl_2_ versus 0.59M NaCl – the precipitates formed in CaCl_2_ redissolve in NaCl solution, whereas in MgCl_2_ solution the mucin aggregates undergo liquid-solid phase separation. This suggests the formation of ion bridges between Mg^2+^ ions and mucin backbone.

## Conclusion

In this study, we investigate the phase behavior of mucin aggregates from gastrointestinal mucus in the addition of multivalent salts. A reentrant phase transition is observed with trivalent and tetravalent salts. Mucin aggregates exhibit liquid-solid phase separation in divalent salts such as Ca^2+^, Cu^2+^ and Ba^2+^, whereas the solution remains homogeneous across all tested concentrations of MgCl_2_. The ability of multivalent salts in promoting protein participates follows the order: Fe^3+^>Al^3+^>Ba^2+^>Cu^2+^>Ca^2+^>Mg^2+^, which is different from the Hofmeister series. What’s more, precipitates formed in CaCl_2_ solution can be redissolved through ion exchange by adding NaCl or MgCl_2_. In contrast, precipitates formed in FeCl_3_ do not redissolve even in high salt conditions (4M NaCl or 2M MgCl_2_), implying the formation of coordination complexes between Fe^3+^ ions and carboxylate groups (-COO^-^) on the mucin backbone.

## Materials and Methods

### Sample preparation

The mucin samples were first centrifuged at 3000 rpm for 10 minutes, and the supernatant was then filtered through a 450 nm PVDF filter to obtain a clarified mucin solution for subsequent experiments. The size of the mucin aggregates was characterized by light scattering using this prepared solution. Separately, solutions of various salts were prepared and filtered through a 220 nm filter. These filtered salt solutions were then added to the filtered mucin solution. After 48 hours, the samples were photographed to document phase behavior. For the phase-separated samples, the supernatant was filtered through a 450 nm filter for light scattering measurements to obtain the size of the mucin aggregates in the presence of different salts.

### Dynamic light scattering (DLS) measurement

The DLS tubes were washed with pure water and acetone separately more than three times. The tubes were dried in the oven overnight, and then we used aluminum foil to wrap them. Distilled acetone through an acetone fountain setup was used to clean up these tubes. The mucin and mucin in various salt solutions were filtered through 450nm PVDF hydrophilic filter into the tubes to remove the dust. The preparation of all the samples should be conducted in a super clean bench. DLS measurement was performed on a commercial spectrometer, which was equipped with a multi-τ digital time correlator (ALV/LSE-5004) using a wavelength of 532nm laser light source. For each sample, the scattering light intensity was correlated at scattering angles 30°, 40°, 50°, 60°, 70° and 90°. The relaxation time was averaged within three samples. Error bars reflect the standard deviation from measurements of three replicate samples. See *SI Appendix* for more details.

### Characterization of scanning electron microscopy (SEM)

Scanning electron microscopy (SEM) images were obtained by a field emission scanning electron microscope (Quanta FEG250) at an operating voltage of 3 kV and 15kV (for samples from BaCl_2_ solution). Before measurements, the precipitates were dispersed on the silicon wafer and dried it at 37°C for 24 hours. The samples were sputtered with an approximately 10 nm thick Au layer using an EM SCD 500 auto fine coater at a current of 20 mA for 2 min.

### Characterization of optical microscope

The precipitate was dispersed on the microscope slide and dried it at 37°C for 1 hour. Bright field images were acquired using a LSM 900 confocal microscope with a 10× Zeiss lens, with the microscope slides were placed in confocal glass bottom dishes.

### Data, Materials, and Software Availability

All data are included in the manuscript and/or *SI Appendix*.

## ACKNOWLEDGEMENTS

This work was supported by the National Key R&D Program of China (Grant No. 2023YFE0124500), the National Natural Science Foundation of China (Grant No. 22273114), the Strategic Priority Research Program of the Chinese Academy of Sciences (Grant No. XDB0770101), and International Partnership Program of the Chinese Academy of Sciences (Grant No. 027GJHZ2022061FN).

